# Tight junction membrane proteins regulate the mechanical resistance of the apical junctional complex

**DOI:** 10.1101/2023.08.02.551232

**Authors:** Thanh Phuong Nguyen, Tetsuhisa Otani, Motosuke Tsutsumi, Sachiko Fujiwara, Tomomi Nemoto, Toshihiko Fujimori, Mikio Furuse

## Abstract

Epithelia must be able to resist mechanical force to preserve tissue integrity. While intercellular junctions are known to be important for the mechanical resistance of epithelia, the roles of tight junctions (TJs) remain to be established. We previously demonstrated that epithelial cells devoid of the TJ membrane proteins claudins and JAM-A completely lack TJs and exhibit focal breakages of their apical junctions. Here, we demonstrate that apical junctions undergo spontaneous fracture when claudin/JAM-A-deficient cells are exposed to mechanical stress. The junction fracture was accompanied by actin disorganization, and actin polymerization was required for apical junction integrity in the claudin/JAM-A-deficient cells. Further deletion of CAR resulted in the disruption of ZO-1 molecule ordering at cell junctions, accompanied by severe defects in apical junction integrity. These results demonstrate that TJ membrane proteins regulate the mechanical resistance of the apical junctional complex in epithelial cells.

**Summary:** Tight junction membrane proteins claudins, JAM, and CAR coordinately regulate the nanometer-scale organization of ZO-1 molecules, and are required for the mechanical resistance of apical junctions in epithelial cells.

## Introduction

Epithelia cover the body and act as barriers to segregate the internal body from the external environment. Epithelial cells adhere to one another via intercellular junctions that consist of tight junctions (TJs), adherens junctions (AJs), and desmosomes, collectively known as the apical junctional complex (Farquhar and Palade, 1963). TJs are located at the most apical region of the intercellular junctions where adjacent plasma membranes are closely apposed and restrict free diffusion of solutes across the paracellular space (Farquhar and Palade, 1963; Anderson and van Itallie, 2009; Shen et al., 2011; Zihni et al., 2016; Otani and Furuse, 2020).

TJs are composed of integral membrane proteins, including claudins, TJ-associated MARVEL domain-containing proteins (TAMPs: occludin, tricellulin, MarvelD3), and immunoglobulin superfamily proteins (e.g. junctional adhesion molecules [JAMs], and coxsackie and adenovirus receptor [CAR]) (Furuse et al., 1993; Furuse et al., 1998; Martin-Padura et al., 1998; Cohen et al., 2001; Ikenouchi et al., 2005; Steed et al., 2009; Raleigh et al., 2010). These membrane proteins interact with ZO family scaffolding proteins through their cytoplasmic region. ZO family proteins are multidomain scaffolding proteins required for TJ formation (Umeda et al., 2006; Phua et al., 2014; Otani et al., 2019; Rouaud et al, 2020), and have N-terminal PDZ domains that interact with claudins and JAMs (Itoh et al., 1999; Bazzoni et al., 2000; Ebnet et al., 2000; Itoh et al., 2001) and a C-terminal actin-binding region involved in the epithelial barrier function (Fanning et al., 1998; Fanning et al., 2002; Belardi et al., 2020).

Epithelia are subjected to various mechanical stresses, including morphogenetic movements, visceral muscle contractions, cytokinesis, and cell death within the epithelial sheet. The intercellular junctions are involved in maintaining the epithelial integrity under these types of stress. Loss of AJs results in disruption of tissue integrity (Matsunaga et al., 1988; Takeichi, 1991; Kintner, 1992; Harris et al., 2012), while dysfunction of desmosomes leads to pemphigus, which is characterized by skin blistering (Amagai et al., 1991). Recent findings have suggested that mechanosensor molecules are localized at the intercellular junctions and participate in mechanotransduction (Charras and Yap, 2018; Angulo-Urarte et al., 2020). For example, α-catenin, which acts as a linker between the cadherin-catenin complex and actin filaments at AJs, adopts a closed conformation that unfolds in response to tension to unmask a cryptic binding site for vinculin (Yonemura et al., 2010). Similarly, ZO-1 adopts a folded conformation that unfolds in response to application of mechanical force, allowing recruitment of the transcription factor DbpA to TJs (Spadaro et al., 2017). However, unlike AJs and desmosomes, the roles of TJs in regulating the mechanical resistance of epithelia have remained unclear, partly due to the lack of methods to specifically and completely perturb TJs in epithelial cells.

We have recently performed systematic genome editing studies on TJ proteins, and demonstrated that MDCK II cells deficient in claudins (claudin-2/4/3/7/1) and JAM-A (claudin/JAM-A knockout [KO] cells) exhibit specific and complete loss of the TJ structure and function (Otani et al., 2019). Although AJs were present in the claudin/JAM-A KO cells, a characteristic focal junction breakage phenotype was observed (Otani et al., 2019). However, the underlying mechanism for the phenotype remained unclear.

In the present study, we utilized time-lapse imaging to examine how the junction breakage phenotype arises in claudin/JAM-A KO cells. We found that the apical junctions in the cells showed weak resistance to mechanical stress. Moreover, TJ membrane proteins were required for nanometer-scale ordering of ZO-1 molecules, which appeared to be critical for the resilience of apical junctions against mechanical stress. The present findings establish a role for TJs in the mechanical resistance of epithelial cell junctions.

## Results

### Focal apical junctional complex defects are observed in claudin/JAM-A KO cells

To characterize the junction breakage phenotype in claudin/JAM-A KO cells, we performed immunofluorescence staining of various cell-cell junction markers. Control MDCK II cells showed continuous ZO-1 staining along the apical junctions (Fig. 1A). In contrast, sporadic discontinuity in ZO-1 staining was observed in claudin/JAM-A KO cells (Fig. 1B’, arrow), and in extreme cases, large gaps in ZO-1 staining were observed (Fig. 1B”, asterisk). The phenotype was consistently observed in multiple clones of claudin/JAM-A KO cells (data not shown). Quantification of the number of cell junction endpoints per unit area confirmed the presence of junction breakage in claudin/JAM-A KO cells, but not in MDCK II, JAM-A KO, or claudin KO cells (Fig. 1C). ZO-2 and occludin staining showed similar defects to ZO-1 staining (Fig. 1D, E). Moreover, AJ proteins including afadin and E-cadherin exhibited discontinuous staining at the apical junctions in claudin/JAM-A KO cells (Fig. 1F–I), while continuous staining of E-cadherin was maintained at lateral cell-cell contacts (Fig. 1J, arrow). These results suggest that claudins and JAM-A are required for the integrity of the apical junctional complex.

**Figure 1.**
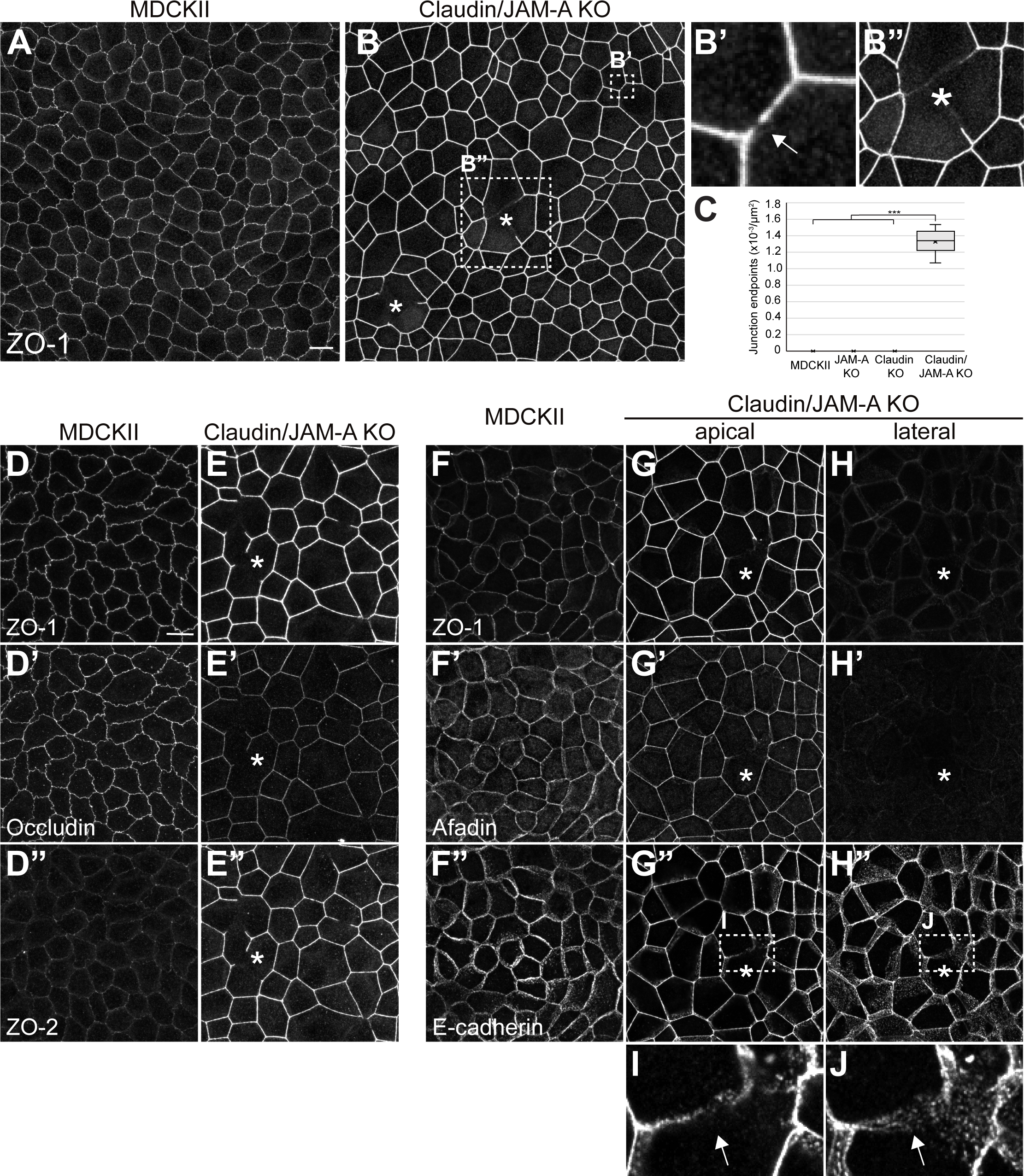
Focal breakage of apical junctions in claudin/JAM-A KO cells. (A) ZO-1 staining of MDCK II cells. A continuous chicken-wire pattern was observed. (B) ZO-1 staining of claudin/JAM-A KO cells. Focal junction breakage was evident, wherein ZO-1 staining was discontinuous (B’, arrow) or large gaps in the ZO-1 network were observed (B”, asterisk). (C) Quantification of the junction breakage phenotype in claudin/JAM-A KO cells. The data represent the numbers of endpoints of ZO-1 staining per unit area and are presented as mean ± SD (*n*=9). ***p<0.0005, compared by *t*-test. (D) ZO-1/occludin/ZO-2 triple staining of MDCK II cells. A continuous chicken-wire pattern was observed for all markers. (E) ZO-1/occludin/ZO-2 triple staining of claudin/JAM-A KO cells. Junction breakage (asterisks) was observed for all markers. (F) ZO-1/afadin/E-cadherin triple staining of MDCK II cells. A continuous network of ZO-1, afadin, and E-cadherin was evident in MDCK II cells. (G–J) ZO-1/afadin/E-cadherin triple staining of claudin/JAM-A KO cells. Junction breakage was observed for ZO-1 (G, asterisk) and afadin (G’, asterisk). Discontinuity of E-cadherin staining was observed at the apical AJs (G”, asterisk; I, arrow), while continuous E-cadherin staining was maintained at the lateral cell contacts (H”, J, arrow; asterisk indicates the region with junction breakage). Maximum intensity projections of apical (G, I) and lateral (H, J) confocal sections are shown. Scale bars: 10 μm.

### Mechanical stress triggers junction breakage in claudin/JAM-A KO cells

To understand how the junction breakage phenotype arises in claudin/JAM-A KO cells, we stably expressed ZO-1-GFP in control MDCK II cells and claudin/JAM-A KO cells and performed time-lapse imaging. Multiple clones were established for both cell lines, and representative data are shown. In control MDCK II cells, the continuity of ZO-1-GFP was maintained during cell stretching (Fig. 2A; Video 1) and cytokinesis (Fig. 2B; Video 1), consistent with previous reports (Jinguji and Ishikawa, 1992; Higashi et al., 2016). These results indicate that the cell junctions in control cells were robust against mechanical stress. Meanwhile, although the chicken-wire pattern of ZO-1-GFP was established in claudin/JAM-A KO cells during monolayer formation, the junctions underwent breakage upon spontaneous stretching of the cells (Fig. 2C; Video 2). The broken junctions were reassembled within a few hours, implying that the junction repair pathways (Stephenson et al., 2019; Higashi et al., 2022) were not severely compromised in the claudin/JAM-A KO cells. Similar junction breakage was observed when claudin/JAM-A KO cells underwent cytokinesis (Fig. 2D; Video 2), suggesting that the junction breakage occurred when the cell junctions were subjected to mechanical stress. Quantification of the cell circumference and the fluorescence intensity of ZO-1-GFP confirmed that the junction breakage occurred upon acute cell stretching (Fig. 2E, F). Measurements from multiple stretching events revealed that the cell junctions in claudin/JAM-A KO cells underwent breakage above a certain threshold (∼30% stretching), while the cell junctions in control MDCK II cells withstood a similar degree of stretching (Fig. 2G). This observation of a stretching threshold strongly suggests that the junction breakage in claudin/JAM-A KO cells involves a mechanical fracture of the cell junctions. Taken together, these results indicate that claudins and JAM-A regulate the mechanical resistance of apical junctions.

**Figure 2.**
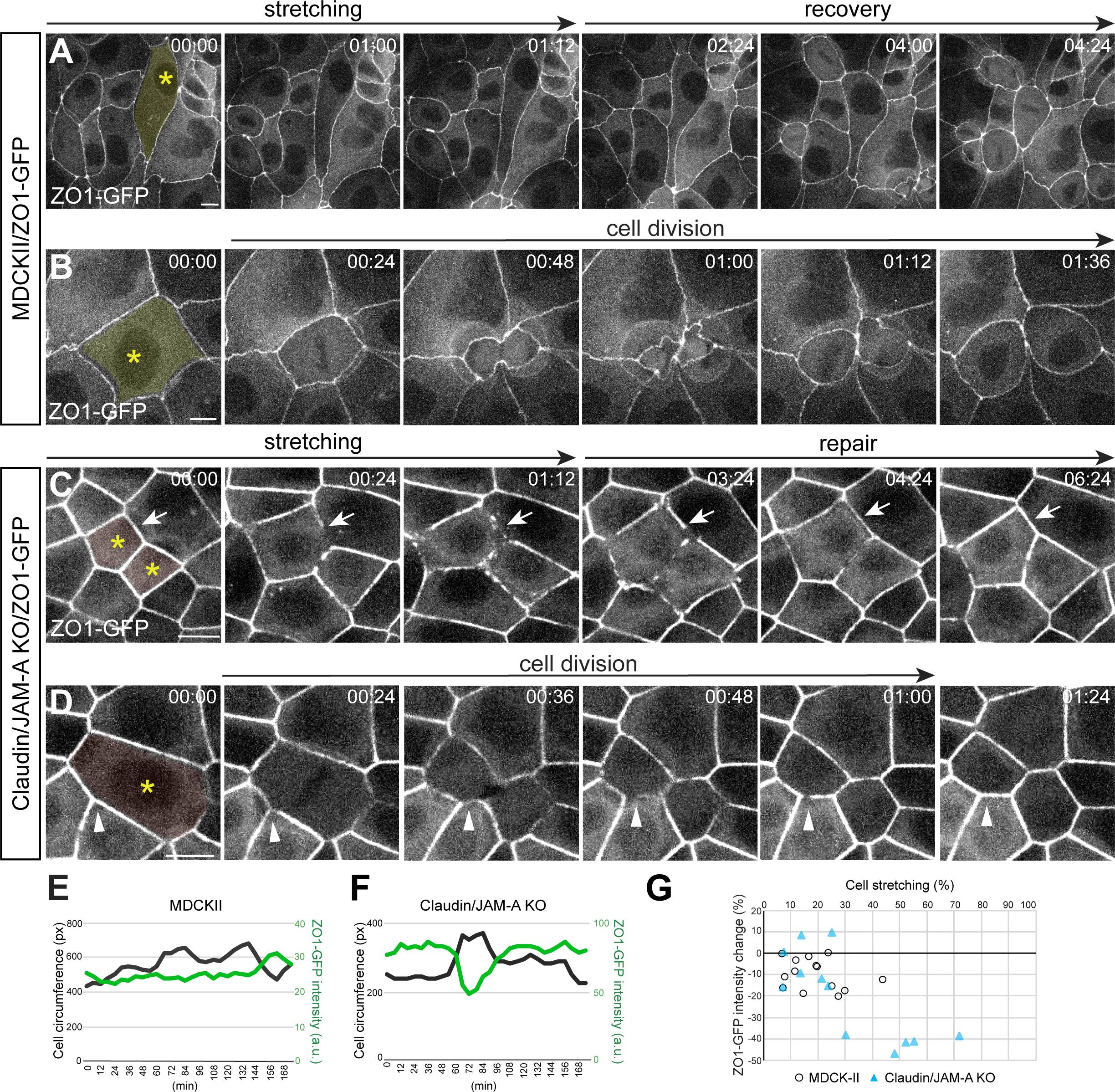
Mechanical stress triggers junction breakage in claudin/JAM-A KO cells. (A) Snapshots from a movie of MDCK II cells expressing ZO-1-GFP. ZO-1-GFP maintained its continuity while the cells underwent stretching. (B) Snapshots from a movie of MDCK II cells expressing ZO-1-GFP, wherein a cell underwent cytokinesis. The junction continuity was maintained when cells underwent stretching during the cytokinesis. (C) Snapshots from a movie of claudin/JAM-A KO cells expressing ZO-1-GFP. ZO-1-GFP became discontinuous or fragmented when a cell underwent acute stretching. The junctions were subsequently reassembled within 6 h. (D) Snapshots from a video of claudin/JAM-A KO cells expressing ZO-1-GFP. When a cell underwent cytokinesis, the junctions of the neighboring cells became discontinuous or fragmented. The junctions were reassembled within 2 h. (E) Quantification of the cell circumference (black) and ZO-1 intensity (green) during the cell stretching event in MDCK II cells shown in (A). The ZO-1 intensity did not change significantly during the cell stretching. (F) Quantification of the cell circumference (black) and ZO-1 intensity (green) during the cell stretching event in claudin/JAM-A KO cells shown in (C). The ZO-1 fluorescence intensity showed an acute reduction upon cell stretching. (G) Scatter plot of the cell stretching degrees and the ZO-1 intensity changes in multiple cell stretching events in MDCK II cells (white circles) and claudin/JAM-A KO cells (blue triangles). In MDCK II cells, the ZO-1 intensity showed a mild reduction during cell stretching, which could have arisen from dilution of ZO-1 as the cell circumference increased. In claudin/JAM-A KO cells, the ZO-1 intensity did not change markedly when the cells underwent <30% stretching. However, a large reduction in ZO-1 intensity was observed when the cells were stretched by >30%, corresponding to the junction breakage events. Time scales: h: min. Scale bars: 10 μm.

### The actin cytoskeleton is disorganized at junction breakage sites

Because the actin cytoskeleton plays a central role in regulating the apical junction integrity, we focused on the organization of the actin cytoskeleton during junction breakage. In control MDCK II cells, actin filaments visualized by phalloidin and myosin IIA and IIB were weakly localized at the cell-cell junctions with additional staining in the apical cytoplasm (Fig. S1A–E). In contrast, F-actin and myosin IIA and IIB were strongly localized in the extensively developed circumferential actin bundles underlying the apical junctional complex in claudin/JAM-A KO cells (Fig. S1A’–E’). This localization resulted in the application of strong tension to AJs, as revealed by increased staining of the α-18 antibody, which recognizes an epitope exposed in α-catenin in its extended conformation (Fig. S1F) (Yonemura et al., 2010), although the amount of total α-catenin at the cell-cell junctions was not significantly changed (Fig. S1G). Vinculin, which is recruited to the extended α-catenin (Yonemura et al., 2010), was also strongly accumulated in claudin/JAM-A KO cells (Fig. S1I), indicating that the cell-cell junctions were subjected to increased tension. Both α-18 and vinculin were reduced in the large gaps of ZO-1, confirming the disorganization of AJs at the junction breakage sites (Fig. S1F–J).

Super-resolution imaging using stimulated-emission depletion (STED) revealed that the circumferential actin bundles were disorganized at the junction breakage sites in claudin/JAM-A KO cells. Although the circumferential actin bundles were coalesced and tightly associated with the intact cell-cell junctions in claudin/JAM-A KO cells (Fig. 3A, B), the bundles became loosened at the junction breakage sites (Fig. 3C–E, arrows). The actin bundles remained tightly attached to the cell-cell junctions where focal ZO-1 localization persisted (Fig. 3D, E, arrowheads). Similar results were observed for myosin IIA (Fig. 3F–J) and myosin IIB (Fig. 3K–O). Furthermore, the actomyosin bundles from the adjacent cells were often separated from one another in the large gaps of ZO-1 staining (Fig. 3C, H, M, arrows). These results demonstrate that the junction breakages in claudin/JAM-A KO cells are accompanied by disorganization of the actin cytoskeleton.

**Figure 3.**
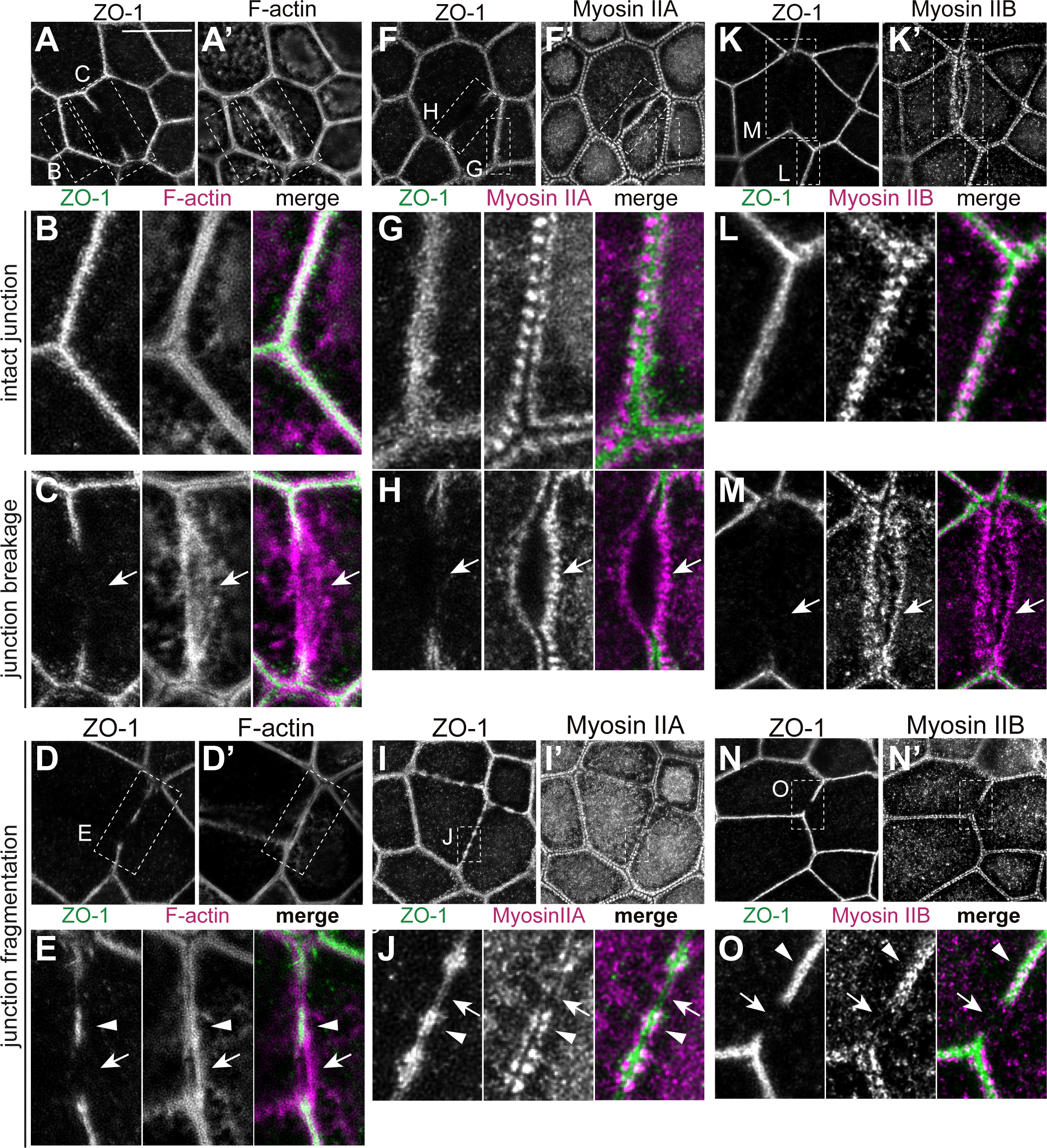
The actin cytoskeleton is disorganized at junction breakage sites in claudin/JAM-A KO cells. The actomyosin organization at the junction breakage sites was observed by STED microscopy. (A–E) F-actin organization at junction breakage sites. The F-actin bundles were loosened at junction breakage sites (arrows), but coalesced and became attached to the cell junctions where ZO-1 localization persisted (arrowheads). (F–J) Myosin IIA localization at junction breakage sites. The myosin IIA bundles were loosened at large junction breakage sites (F–H, arrows). The myosin IIA localization was fragmented at small ZO-1 fragmentation sites (I; J, arrows). Remnant myosin IIA staining was observed at the sites where ZO-1 localization persisted (J, arrowheads). (K–O) Myosin IIB localization at junction breakage sites. The myosin IIB bundles were loosened at large junction breakage sites (K–M, arrows). The myosin IIB localization was fragmented at small ZO-1 fragmentation sites (N; O, arrows). Remnant myosin IIB staining was observed at the sites where ZO-1 localization persisted (O, arrowheads). Scale bar: 10 μm.

We performed two-color time-lapse imaging to examine the spatiotemporal relationship between the actin organization and junction breakage by stably expressing ZO-1-GFP and LifeAct-mCherry in claudin/JAM-A KO cells. The time-lapse imaging revealed that the loosening of the junction-associated actin bundles occurred simultaneously with the breakage of the junctions, suggesting that the junction breakage was tightly correlated with the disorganization of the actin cytoskeleton (Fig. 4A, B; Video 3). Interestingly, the reassembly of the actin bundles preceded the re-localization of ZO-1-GFP during the junction repair process (Fig. 4C, D; Video 4). These results suggest that the actin cytoskeleton plays an important role in regulating the junction integrity.

**Figure 4.**
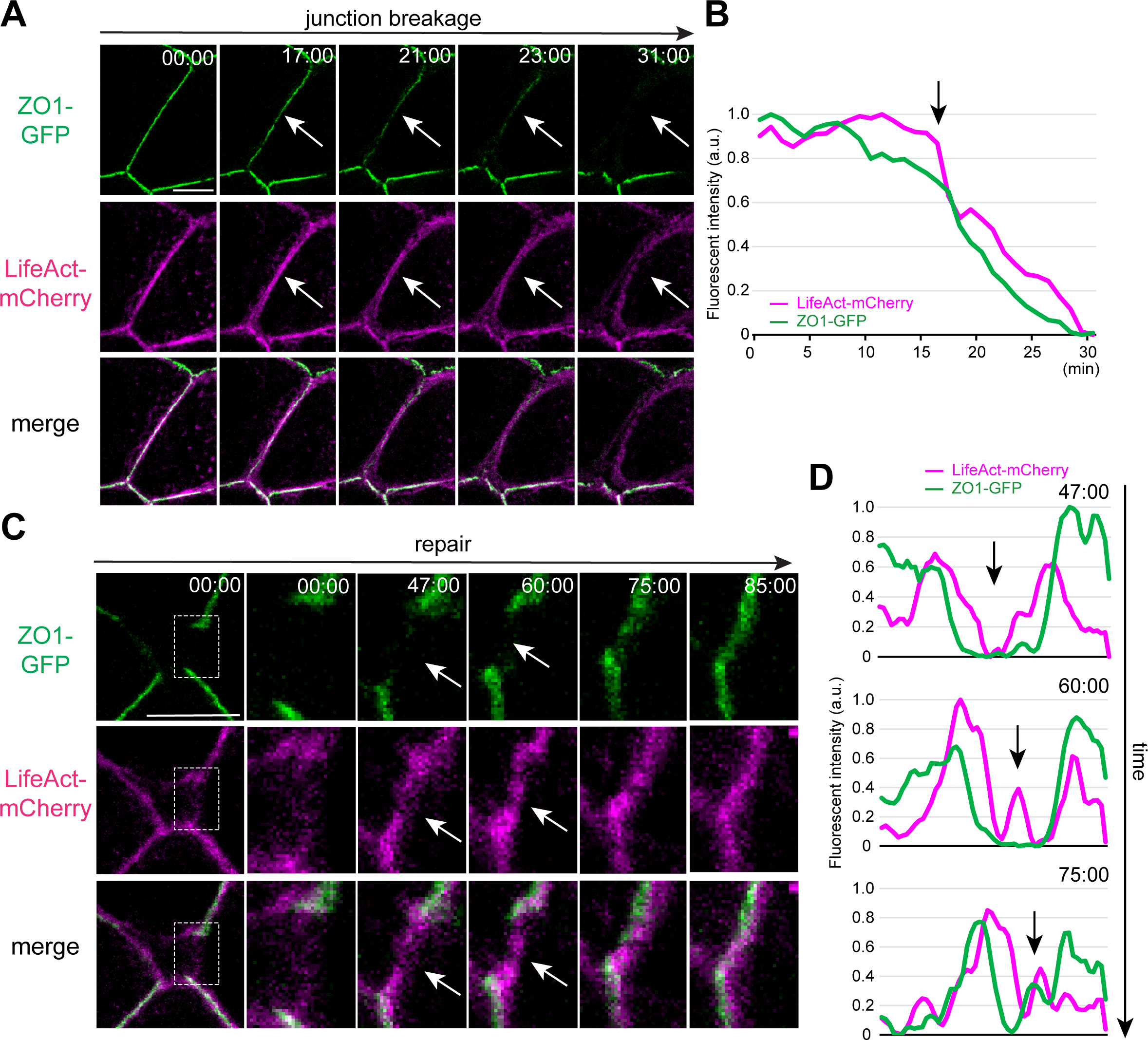
Actin cytoskeleton disorganization accompanies junction breakage in claudin/JAM-A KO cells. (A) Snapshots from a two-color time-lapse movie of claudin/JAM-A KO cells expressing ZO-1-GFP (green) and LifeAct-mCherry (magenta) showing a junction breakage event. The actin bundles were loosened and became detached from the cell junctions during the junction breakage (arrows). (B) Quantification of the junctional fluorescence intensities of ZO-1-GFP (green) and LifeAct-mCherry (magenta) during the junction breakage. The fluorescence intensities of ZO-1-GFP (green) and LifeAct-mCherry (magenta) were simultaneously reduced. The arrow indicates the start of the junction breakage event. (C) Snapshots from a two-color time-lapse movie of claudin/JAM-A KO cells expressing ZO-1-GFP (green) and LifeAct-mCherry (magenta) showing the junction recovery process after junction breakage. The boxed regions are enlarged in the right panels. The reassembly of the cell junction-associated actin bundles preceded the recovery of ZO-1-GFP junction accumulation (arrows). (D) Quantification of the junctional fluorescence intensities of ZO-1-GFP (green) and LifeAct-mCherry (magenta) during the junction recovery. The fluorescence intensity of LifeAct-mCherry (magenta) recovered before the fluorescence intensity of ZO-1-GFP (green) at the junction breakage site (arrows). Time scales: min: s. Scale bars: 10 μm.

### Actin polymerization regulates the apical junction integrity in claudin/JAM-A KO cells

To investigate whether the disorganization of the actin cytoskeleton has a causal role in regulating the junction integrity, we pharmacologically perturbed the actomyosin cytoskeleton. The RhoA-ROCK signaling pathway plays a central role in regulating the organization of the actomyosin cytoskeleton. The ROCK inhibitor Y-27632 did not affect the continuity of ZO-1 staining in control MDCK II cells (Fig. 5A, B) but induced an exaggerated junction breakage phenotype in claudin/JAM-A KO cells (Fig. 5B’, F). ROCK was reported to stimulate myosin II-dependent contraction through phosphorylation of the myosin II regulatory light chain (Amano et al., 1996; Kimura et al., 1996), and to promote actin polymerization by inhibiting cofilin via LIMK (Maekawa et al., 1999; Yang et al., 1998). These observations prompted us to dissect the downstream signaling pathways of ROCK. The myosin II inhibitor blebbistatin did not modify the junction breakage phenotype in claudin/JAM-A KO cells (Fig. 5C’, F). However, inhibition of LIMK by BMS-5 induced discontinuity of the ZO-1 staining in claudin/JAM-A KO cells (Fig. 5D’, F), suggesting that ROCK-LIMK-cofilin dependent regulation of actin polymerization was important for the cell junction integrity.

**Figure 5.**
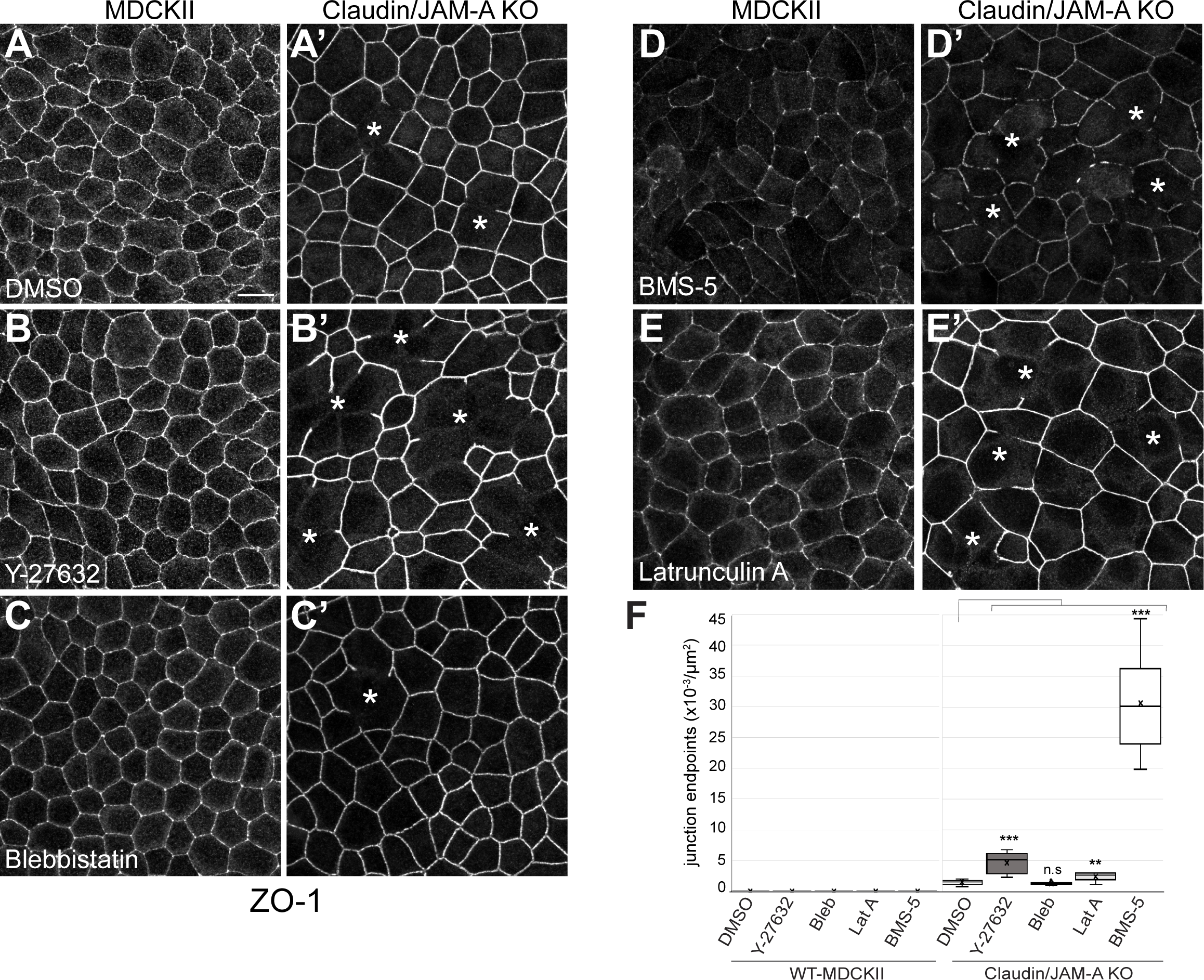
Actin polymerization is required for the apical junction integrity in claudin/JAM-A KO cells. (A) ZO-1 staining of MDCK II cells treated with DMSO. ZO-1 formed a continuous chicken-wire pattern. (A’) ZO-1 staining of claudin/JAM-A KO cells treated with DMSO. Junction breakage was occasionally observed (asterisks). (B) ZO-1 staining of MDCK II cells treated with ROCK inhibitor Y-27632 (10 μM, 3 h). ZO-1 staining was continuous. (B’) ZO-1 staining of claudin/JAM-A KO cells treated with ROCK inhibitor Y-27632 (10 μM, 3 h). The junction breakage phenotype was exaggerated (asterisks). (C) ZO-1 staining of MDCK II cells treated with myosin II inhibitor blebbistatin (100 μM, 3 h). ZO-1 staining was continuous. (C’) ZO-1 staining of claudin/JAM-A KO cells treated with myosin II inhibitor blebbistatin (100 μM, 3 h). The junction breakage phenotype (asterisk) was comparable to that in DMSO-treated claudin/JAM-A KO cells. (D) ZO-1 staining of MDCK II cells treated with LIMK inhibitor BMS-5 (10 μM, 3 h). ZO-1 staining was reduced but remained continuous. (D’) ZO-1 staining of claudin/JAM-A KO cells treated with ROCK inhibitor LIMK inhibitor BMS-5 (10 μM, 3 h). Extensive fragmentation of cell junctions was observed (asterisks). (E) ZO-1 staining of MDCK II cells treated with actin polymerization inhibitor latrunculin A (0.3 μM, 1 h). ZO-1 staining remained continuous. (E’) ZO-1 staining of claudin/JAM-A KO cells treated with actin polymerization inhibitor latrunculin A (0.3 μM, 1 h). Extensive fragmentation of cell junctions was observed (asterisks). (G) Quantification of the junction breakage phenotype. Data represent the numbers of endpoints of ZO-1 staining per unit area and are presented as mean ± SD (*n*=9). **p<0.005, ***p<0.0005, compared by *t*-test. Scale bar: 10 μm.

Consistently, latrunculin A, which inhibits actin polymerization, enhanced the junction breakage phenotype in claudin/JAM-A KO cells (Fig. 5E’, F) at concentrations that did not severely affect the junction integrity in control MDCK II cells (Fig. 5E, F). These results suggest that actin polymerization is required for the apical junctional complex integrity in claudin/JAM-A KO cells.

### Both the *trans*-interaction and ZO-1 binding of claudins and JAM-A are important for the mechanical resistance of cell junctions

The above results suggested that claudins and JAM-A together with the actin cytoskeleton regulate the mechanical resistance of the apical junctional complex. To further gain insight into how claudins and JAM-A regulate the junction integrity, we performed structure-function analyses of claudins and JAM-A using a mutant that inhibits the *trans*-interaction and strand formation of claudins (claudin-1[F147A]) (Piontek et al., 2008; Suzuki et al., 2014), a mutant that inhibits the *cis*-dimerization and *trans*-interaction of JAM-A (JAM-A[ΔDL1]) (Monteiro et al., 2014), and mutants that uncouple ZO-1 from claudins (claudin-1[ΔYV]) and JAM-A (JAM-A[ΔLV]) (Fig. 6A). The inability of claudin-1[F147A] and JAM-A[ΔDL1] to *trans*-interact (Fig. S2A–F), and the inability of claudin-1[ΔYV] and JAM-A[ΔLV] to recruit ZO-1 to cell-cell contacts (Fig. S2G–R) were confirmed by expressing these molecules in L fibroblasts. Full-length claudin-1 and JAM-A and their mutants were stably expressed in claudin/JAM-A KO cells, and their ability to rescue the junction breakage phenotype was assessed. Full-length claudin-1 and JAM-A completely rescued the junction breakage phenotype in claudin/JAM-A KO cells (Fig. 6B, E, H). In contrast, neither claudin-1[F147A] nor claudin-1[ΔYV] rescued the junction breakage phenotype, and large gaps in ZO-1 staining were observed (Fig. 6C, D, H). JAM-A[ΔLV] also failed to rescue the junction breakage phenotype (Fig. 6G, H). JAM-A[ΔDL1] suppressed the formation of large gaps, but focal discontinuity in ZO-1 staining was frequently observed (Fig. 6F, H). These results suggest that both the *trans*-interaction and the ability to interact with ZO-1 are important for the mechanical resistance of apical cell junctions.

**Figure 6.**
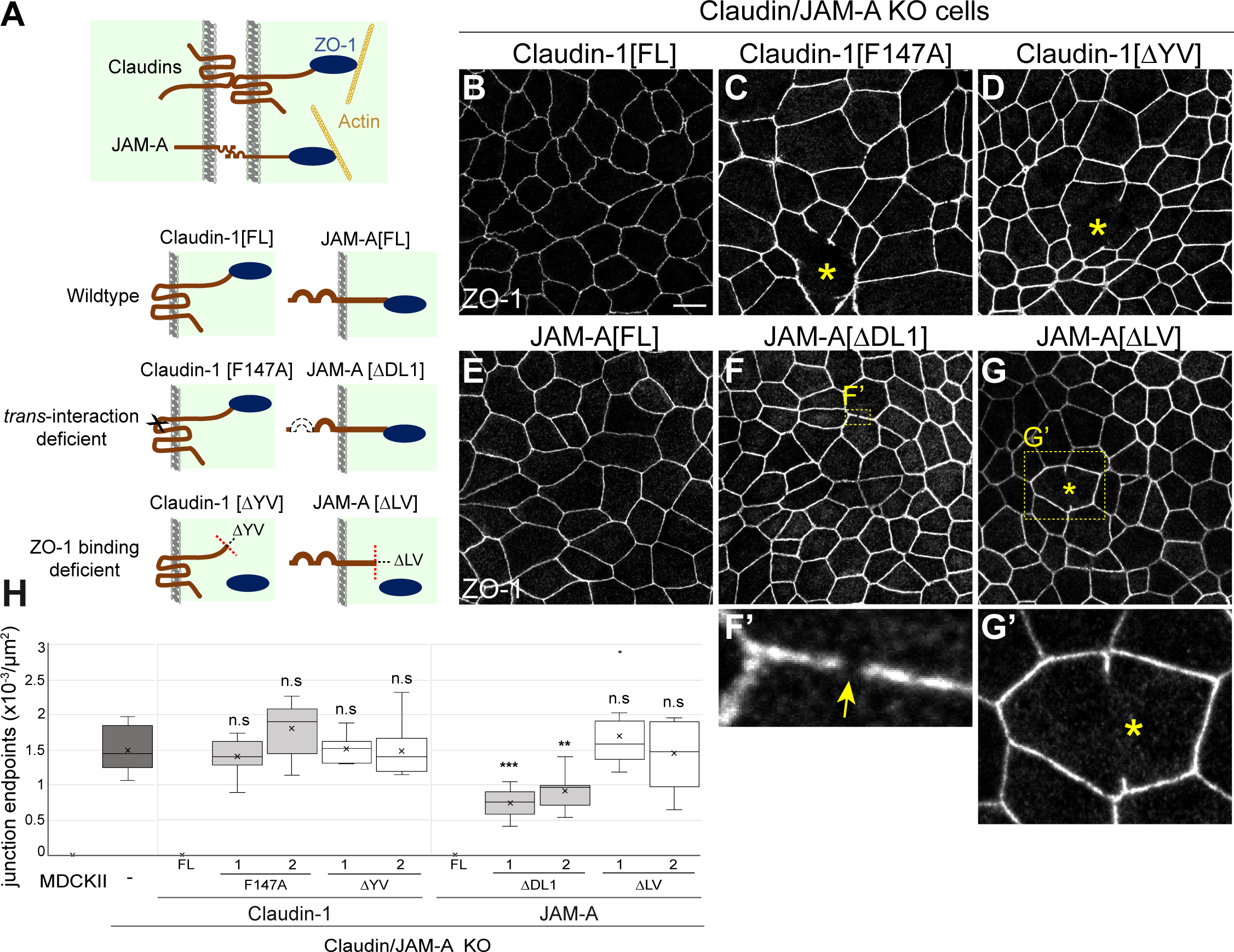
Both the *trans*-interaction and ZO-1 binding of claudin and JAM-A are important to support apical junction integrity. (A) Schematic illustration of the constructs used in the study. Claudin-1[F147A] is a mutant that inhibits the *trans*-interaction and strand formation, while claudin-1[ΔYV] lacks the PDZ-binding motif required for the interaction with ZO-1. JAM-A[ΔDL1] inhibits the *cis*-dimerization and *trans*-interaction, while JAM-A[ΔLV] lacks the PDZ-binding motif required for the interaction with ZO-1. (B–D) ZO-1 staining of claudin/JAM-A KO cells rescued with claudin-1 full-length (B), claudin-1[F147A] (C), or claudin-1[ΔYV] (D). While claudin-1 full-length completely restored the junction continuity (B), claudin-1[F147A] (C) and claudin-1[ΔYV] (D) failed to rescue the junction breakage phenotype (asterisks). (E–G) ZO-1 staining of claudin/JAM-A KO cells rescued with JAM-A full-length (E), JAM-A[ΔDL1] (F), or JAM-A[ΔLV] (G). JAM-A full-length completely rescued the junction breakage phenotype (E). JAM-A[ΔDL1] suppressed the formation of large gaps but focal discontinuity of cell junctions was frequently observed (F’, arrow). JAM-A[ΔLV] failed to rescue the junction breakage phenotype (G’, asterisks). (H) Quantification of the junction breakage phenotype. Data represent the numbers of endpoints of ZO-1 staining per unit area and are presented as mean ± SD (*n*=9). **p<0.005, ***p<0.0005, compared with claudin/JAM-A KO cells by *t*-test. Scale bar: 10 μm.

### Claudins and JAM-A regulate the conformation of ZO-1

The structure-function analyses of claudins and JAM-A suggested that the transmembrane linkage of neighboring cells via claudin and JAM-A and the actin cytoskeleton through ZO-1 was important for the mechanical resistance of cell junctions. Recent reports have indicated that ZO-1 may act as a mechanosensor and undergoes a conformational change in response to tension (Spadaro et al., 2017). These findings raise the possibility that claudins and JAM-A may regulate the mechanical resistance of the apical cell junctions by regulating the conformational status of ZO-1.

To investigate this possibility, we examined the molecular conformation of ZO-1 by adding distinct epitope tags to its N- and C-termini and measuring the distance between the termini using super-resolution STED microscopy (Fig. 7A). MDCK II cells or claudin/JAM-A KO cells stably expressing ZO-1 with an N-terminal HA tag and a C-terminal Flag tag were cocultured with non-transfected cells, and the junctions between the transfected and non-transfected cells were observed (Fig. 7A). In control MDCK II cells, the STED imaging revealed that the N-terminal HA tag was localized proximal to the membrane, while the C-terminal Flag tag was located more distally on the cytoplasmic side (Fig. 7B). Quantitative measurement of the distance between the N- and C-termini showed that 73% of the ZO-1 molecules had >60-nm distance between the termini (Fig. 7D), suggesting that ZO-1 had adopted an open conformation as reported previously (Spadaro et al., 2017). In claudin/JAM-A KO cells, the oriented localization of ZO-1 was maintained, with the N-terminus localized at the membrane-proximal region and the C-terminus located on the cytoplasmic side (Fig. 7C). However, quantitative analysis revealed that the distance between the N- and C-termini of ZO-1 was shorter in claudin/JAM-A KO cells, and only 53% of the ZO-1 molecules in claudin/JAM-A KO cells had >60-nm distance between the termini (Fig. 7D), indicating that ZO-1 preferred a closed conformation in claudin/JAM-A KO cells. These results suggest that claudins and JAM-A regulate the conformation of ZO-1 molecules.

**Figure 7.**
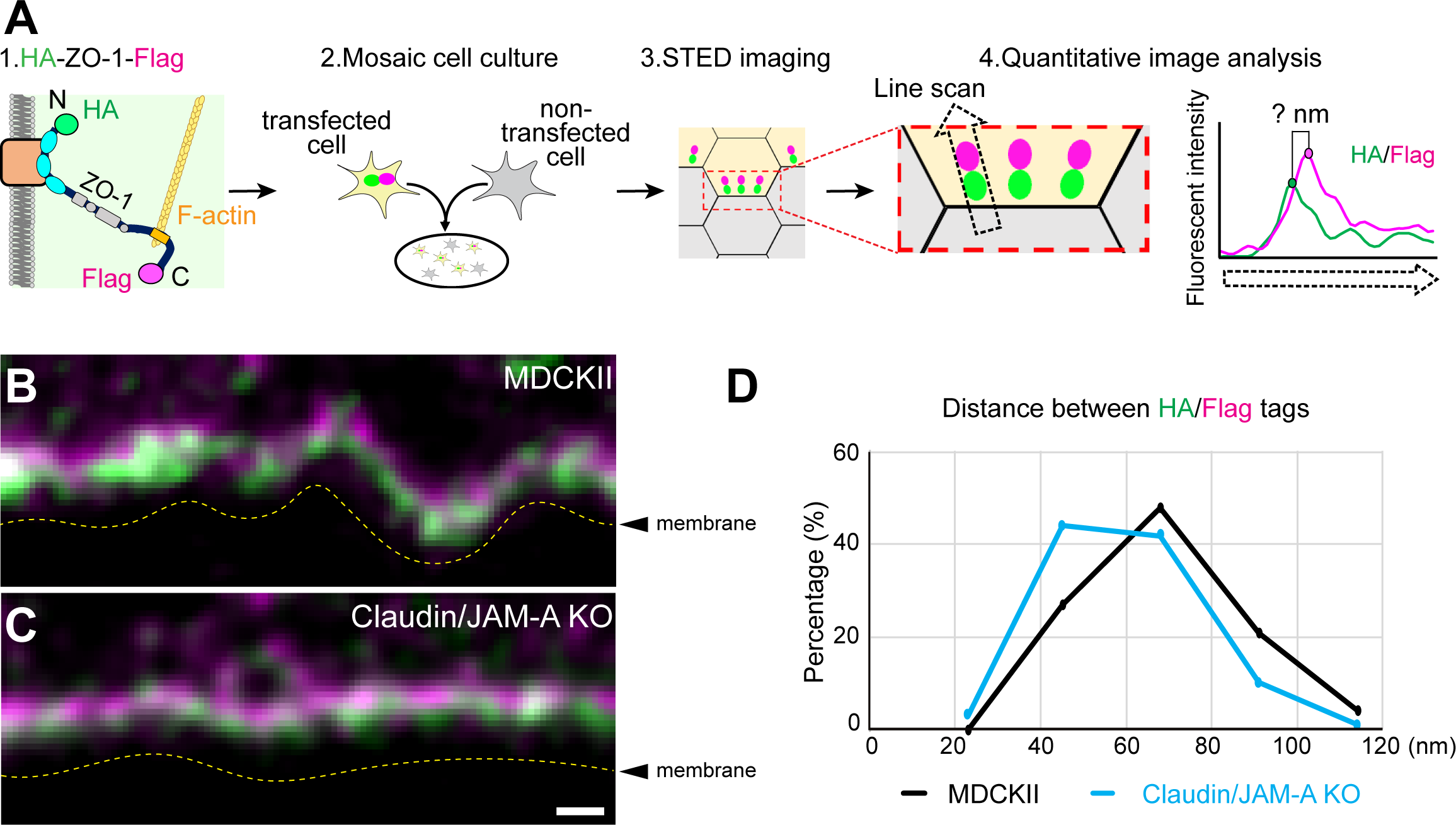
Claudins and JAM-A regulate the conformation of ZO-1. (A) Schematic illustration of the experimental design. MDCK II cells or claudin/JAM-A KO cells stably expressing ZO-1 with an N-terminal HA tag (green) and a C-terminal Flag tag (magenta) were cocultured with non-transfected cells, and the junctions between the transfected and the non-transfected cells were observed. A line scan was performed across paired signals of the HA/Flag tag puncta, and the distance between the HA and Flag tags was measured. (B) STED observation of double staining for the N-terminal HA tag (green) and C-terminal Flag tag (magenta) of HA-ZO-1-Flag expressed in MDCK II cells. The N-terminal HA tag was localized proximal to the membrane, while the C-terminal Flag tag was located more distally on the cytoplasmic side. (C) STED observation of double staining for the N-terminal HA tag (green) and C-terminal Flag tag (magenta) of HA-ZO-1-Flag expressed in claudin/JAM-A KO cells. The N-terminus localized at the membrane-proximal region, while the C-terminus was located on the cytoplasmic side. (D) Quantification of the distance between the N-terminal HA tag (green) and C-terminal Flag tag (magenta) in MDCK II cells and claudin/JAM-A KO cells. ZO-1 tended to adopt an elongated conformation (>60 nm) in MDCK II cells, while it preferred a closed conformation (<60 nm) in claudin/JAM-A KO cells (*n*=100 particles from 10–11 junctions). The plasma membrane outlines are indicated by the dotted lines in (B) and (C). Scale bar: 200 nm.

### CAR, claudins, and JAM-A are essential for the nanometer-scale ordering of ZO-1

Although the above results demonstrated that claudins and JAM-A regulate the conformation of ZO-1, the N-terminus of ZO-1 was still located at the membrane-proximal region in claudin/JAM-A KO cells, suggesting that other membrane proteins with PDZ-binding motifs may be involved in anchoring the N-terminal PDZ domains of ZO-1 to the membrane-proximal region in these cells. This prompted us to examine the contributions of occludin and CAR, as integral membrane proteins that are localized at TJs and interact with ZO-1 (Furuse et al., 1993; Furuse et al., 1994, Cohen et al., 2001). For this, we generated claudin/JAM-A/occludin KO cells and claudin/JAM-A/CAR KO cells by CRISPR/Cas9-mediated genome editing and analyzed their phenotypes (Fig. S3). Multiple clones were isolated, and all clones showed identical phenotypes.

ZO-1 with an N-terminal HA tag and a C-terminal Flag tag was stably expressed in claudin/JAM-A/occludin KO cells and claudin/JAM-A/CAR cells, and the conformation of ZO-1 was examined. STED imaging revealed that the molecular orientation of ZO-1 remained unaltered in claudin/JAM-A/occludin KO cells, with the N-terminus localized at the membrane-proximal region and the C-terminus located on the cytoplasmic side (Fig. 8A, B). In contrast, the parallel orientation of ZO-1 molecules was severely perturbed in claudin/JAM-A/CAR KO cells, and the N- and C-termini of ZO-1 were randomly localized within the cell junctions (Fig. 8C). The disordering of ZO-1 molecules in claudin/JAM-A/CAR KO cells was confirmed by quantification of the order parameter (Fig. 8D). These results demonstrate that claudins, JAM-A, and CAR anchor the ZO-1 N-terminus to the membrane-proximal region and regulate the nanometer-scale ordering of ZO-1 molecules.

**Figure 8.**
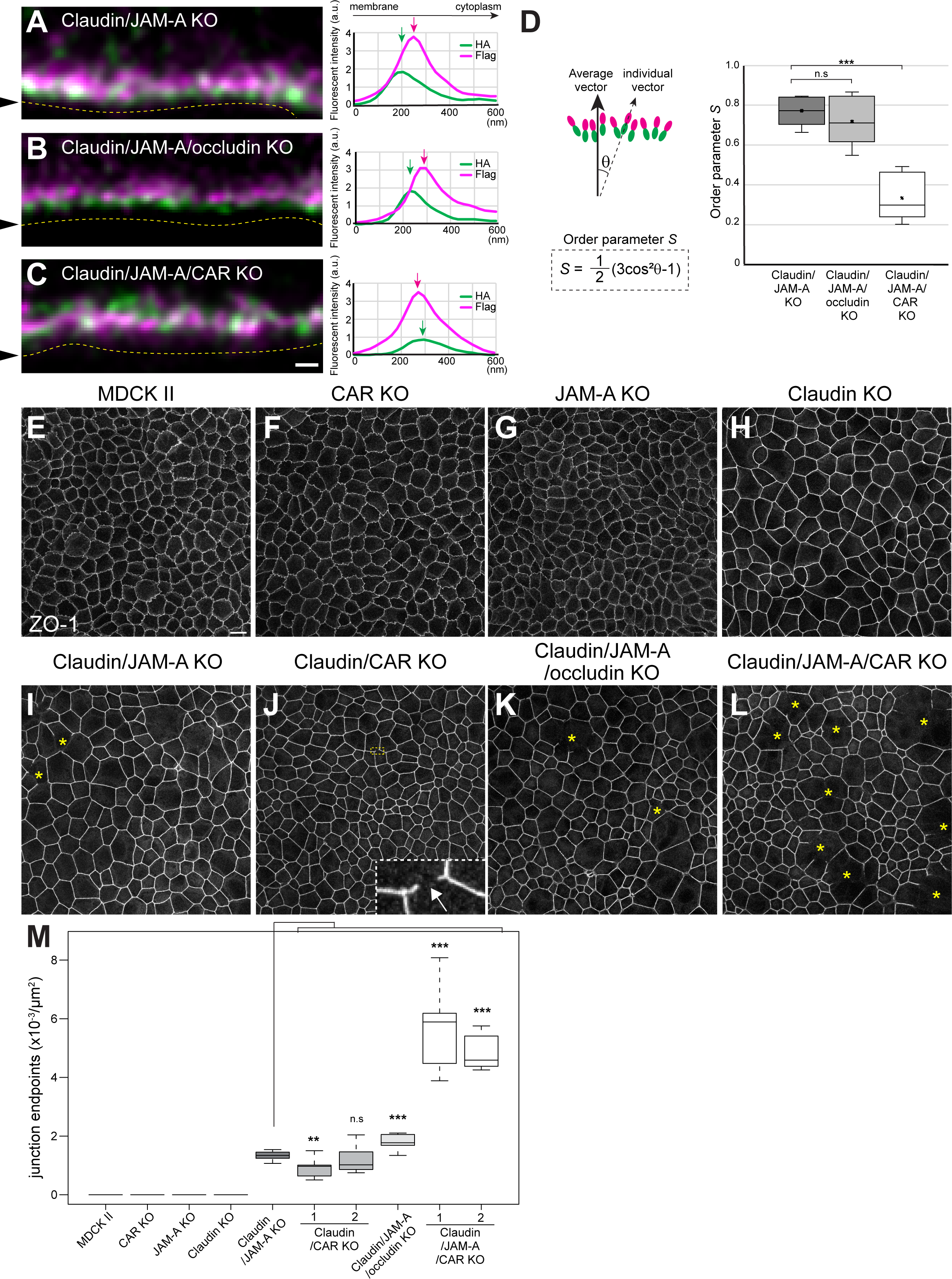
CAR acts in conjunction with claudins and JAM-A to regulate the nanometer-scale ordering of ZO-1 and apical junction integrity. (A) STED observation of double staining for the N-terminal HA tag (green) and C-terminal Flag tag (magenta) of HA-ZO-1-Flag expressed in claudin/JAM-A KO cells. The N-terminus was localized at the membrane-proximal region, while the C-terminus was located on the cytoplasmic side. The line scan shows that the N- and C-termini were separated. (B) STED observation of double staining for the N-terminal HA tag (green) and C-terminal Flag tag (magenta) of HA-ZO-1-Flag expressed in claudin/JAM-A/occludin KO cells. The N-terminus was localized at the membrane-proximal region while the C-terminus was located on the cytoplasmic side. The line scan showed that the N- and C-termini were separated. (C) STED observation of double staining for the N-terminal HA tag (green) and C-terminal Flag tag (magenta) of HA-ZO-1-Flag expressed in claudin/JAM-A/CAR KO cells. The N- and C-termini of ZO-1 were randomly oriented at the membrane-proximal region. The line scan shows that the localization of the N- and C-termini was random. It should be noted that a thick line was used as the region of interest for the line scans shown in (A)–(C), resulting in the averaging of multiple puncta. (D) Quantification of the order parameter *S*. The angle of the HA/Flag pairs was measured, and the deviation angle θ for individual vectors from the average vector was determined. The order parameter *S* was defined as *S* = (3cos^2^θ-1)/2 and showed that the nanometer-scale ordering of ZO-1 molecules at cell junctions was disorganized in claudin/JAM-A/CAR KO cells (*n*=5–6 junctions). (E–L) ZO-1 staining in MDCK II (E), CAR KO (F), JAM-A KO (G), claudin quintuple KO (H), claudin/JAM-A KO (I), claudin/CAR KO (J), claudin/JAM-A/occludin KO (K), and claudin/JAM-A/CAR KO (L) cells. Junction breakage was not observed in MDCK II (E), CAR KO (F), JAM-A KO (G), or claudin KO (H) cells. Claudin/JAM-A KO (I) and claudin/JAM-A/occludin KO (K) cells showed junction breakage with occasional large gaps (asterisks), while claudin/CAR KO cells (J) showed discontinuity in cell junctions (arrow). Claudin/JAM-A/CAR KO cells (L) showed an exaggerated junction breakage phenotype, and numerous large gaps were observed (asterisks). (M) Quantification of the junction breakage phenotype. Data represent the numbers of endpoints of ZO-1 staining per unit area and are presented as mean ± SD (*n*=9). **p<0.005, ***p<0.0005, compared by *t*-test. Scale bars: (A–C) 200 nm; (E–L) 10 μm.

### CAR, claudins, and JAM-A coordinately regulate the apical cell junction integrity

Finally, we examined the junction integrity in claudin/JAM-A/occludin KO and claudin/JAM-A/CAR KO cells. MDCK II, CAR KO, JAM-A KO, and claudin quintuple KO cells did not have the junction breakage phenotype (Fig. 8E–H, M). Claudin/CAR KO cells showed spontaneous discontinuity of ZO-1 staining (Fig. 8J, M, arrow), while claudin/JAM-A/occludin KO cells exhibited a slight enhancement of the junction breakage phenotype compared with claudin/JAM-A KO cells (Fig. 8I, K, M). In contrast, numerous large gaps in ZO-1 staining were observed in claudin/JAM-A/CAR KO cells (Fig. 8L, M, asterisks). Taken together, these results suggest that TJ membrane proteins are required for the nanometer-scale ordering of ZO-1 and the apical cell junction integrity.

## Discussion

Epithelia must be able to resist mechanical stress to maintain the tissue integrity. Although AJs and desmosomes are well known to play important roles in the mechanical resistance of epithelia (Matsunaga et al., 1988; Amagai et al., 1991; Takeichi, 1991; Kintner, 1992; Harris et al., 2012), the roles of TJs remain unclear. In the present study, we utilized claudin/JAM-A KO cells, which exhibit specific and complete disruption of the TJ structure and function, and revealed that TJs are required for the mechanical resistance of the apical junctional complex.

### Roles of TJ membrane proteins in the apical junctional complex organization

AJs and the cadherin-catenin complex have been considered to play central roles in organizing the apical junctional complex, because the perturbation of cadherin-mediated adhesion can severely impair apical junctional complex assembly (Gumbiner and Simons, 1986; Gumbiner et al., 1988; Watabe et al., 1994). However, the roles of TJs in apical junctional complex organization have remained unclear. ZO-1 and ZO-2 have been shown to regulate the organization of AJs and TJs (Umeda et al., 2006; Phua et al., 2014; Otani et al., 2019). It should be noted that although ZO-1 is specifically localized at TJs in mature epithelia, it can also be localized at AJs during junction assembly (Rajasekaran et al., 1996; Ando-Akatsuka et al., 1999), and it remains to be clarified whether the apical junctional complex disorganization in ZO protein-deficient cells reflects the functions of ZO proteins in TJs or AJs. Here, we examined claudin/JAM-A KO cells, in which the TJ structure and function is specifically perturbed, and demonstrated that TJ membrane proteins are essential for the apical junctional complex integrity, despite the presence of AJs and cadherin-mediated adhesion. These findings suggest that, in addition to AJs and the cadherin-catenin complex, TJs play an important role in regulating the apical junctional complex integrity. Because AJs and TJs have distinct mechanosensor molecules (Yonemura et al., 2010; Spadaro et al., 2017), the mechanism for how the AJs and TJs coordinately regulate the mechanical resistance of the apical junctional complex remains an interesting issue to address in future studies.

The present data revealed that TJ membrane proteins regulate the nanometer-scale ordering of ZO-1 molecules. Taken together with previous studies showing essential roles for ZO-1/2 in TJ organization (Umeda et al., 2006; Phua et al., 2014; Otani et al., 2019), the present findings suggest that reciprocal interactions between TJ membrane proteins and ZO proteins are important for the apical junctional complex integrity. It should be noted that ZO proteins still became localized at the apical junctions in the absence of TJ membrane proteins. AJs are present in claudin/JAM-A KO cells (Otani et al., 2019), and ZO-1 can localize to AJs in non-epithelial cells (Itoh et al., 1991; Itoh et al., 1993), suggesting that ZO proteins could be localized to AJs in TJ membrane protein-deficient cells. Other ZO-interacting proteins, including α-catenin and afadin, may play important roles in the recruitment of ZO proteins to the apical junctions in the TJ membrane protein-deficient cells (Rajasekaran et al., 1996; Itoh et al., 1997; Yamamoto et al., 1997; Ooshio et al., 2010). Alternatively, ZO proteins may be recruited to the apical junctions by interacting with F-actin (Itoh et al., 1997).

Another intriguing finding of the present study is the importance of CAR for the apical junctional complex integrity. Although CAR was originally identified as an adenovirus and coxsackievirus receptor (Bergelson et al., 1997), subsequent studies demonstrated that it can also interact with ZO-1 and become localized at TJs (Cohen et al., 2001). It was shown that adenoviruses and coxsackie viruses infect epithelial cells through TJs using CAR as their receptors, and that the epithelial barrier function is transiently disrupted during infection (Walters et al., 2002; Coyne and Bergelson, 2006). Cell culture studies further suggested a role for CAR in the epithelial barrier function (Cohen et al., 2001; Excoffon et al., 2004), and CAR KO mice exhibit complex phenotypes in their epithelial tissues (Pazirandeh et al., 2011). CAR is also expressed in the intercalated discs of the heart and its loss results in cardiac defects (Asher et al., 2005; Dorner et al., 2005; Chen et al., 2006; Lim et al., 2008). These results suggest that CAR functions in parallel with other junctional proteins to coordinately regulate the cell junction organization.

### Role of ZO-1 conformational changes in force resistance

The present results demonstrated that TJ membrane proteins regulate the conformation of ZO-1. Importantly, the nanometer-scale organization of ZO-1 molecules was strongly correlated with the apical junctional complex integrity under mechanical stress, suggesting that the mechanosensitive conformational changes of ZO-1 may play important roles in regulating the mechanical resistance of the apical junctional complex. Because claudins, JAM-A, and CAR interact with the N-terminal PDZ domains in ZO-1 (Itoh et al., 1999; Bazzoni et al., 2000; Ebnet et al., 2000; Itoh et al., 2001), TJ membrane proteins in conjunction with the actomyosin-dependent tensile force applied through the C-terminal actin-binding region of ZO-1 (Fanning et al., 1998; Fanning et al., 2002; Belardi et al., 2020) may regulate the nanometer-scale ordering of ZO-1 molecules at cell junctions. In the TJ membrane protein-deficient cells, the ordering of ZO-1 molecules at cell junctions is compromised due to the absence of its N-terminal binding partners, resulting in a failure to maintain the apical junctional complex integrity under mechanical stress.

The question arises as to how the conformation of ZO-1 can regulate the mechanical resistance of apical junctions. At least three models can be postulated. First, it was reported that ZO-1 undergoes liquid-liquid phase separation, and that this ability of ZO-1 to undergo phase separation is important for TJ formation (Beutel et al., 2019; Schwayer et al., 2019). It was also suggested that the extended conformation of ZO-1 could be a prerequisite for its phase separation (Beutel et al., 2019). These observations suggest that the elongated conformation of ZO-1 may promote ZO-1 condensate formation, which in turn promotes the apical junctional complex integrity. The nanometer-scale alignment of ZO-1 molecules may stabilize the elongated conformation of ZO-1 in a similar manner to polymer brushes, and in the absence of TJ membrane proteins, the extended conformation may become unstable, resulting in a catastrophic collapse of the structure upon application of excessive mechanical stress.

Second, the mechanosensitive conformational change of ZO-1 may act to dissipate excessive mechanical energy, thereby serving as a shock absorber for cell-cell junctions. It was shown that ZO-1 forms multiple intramolecular interactions, and that application of ∼2–4 pN force can disrupt these intramolecular interactions and lead to unfolding and extension of ZO-1 (Spadaro et al., 2017). These findings suggest that when mechanical stress is applied to the apical junctional complex, part of the mechanical energy is used to disrupt the intramolecular interactions of ZO-1, thus dissipating the excessive mechanical energy and protecting the cell junctions. In TJ membrane protein-deficient cells, the force-dependent conformational change of ZO-1 may be abrogated, and when the load exceeds a certain threshold, the apical junctions may undergo mechanical fracture.

Third, ZO-1 was shown to recruit the transcription factor DbpA to TJs in response to its conformational change (Spadaro et al., 2017). DbpA was also shown to regulate epithelial cell proliferation, lumen morphogenesis, and the monolayer integrity of retinal pigment epithelial cells (Balda et al., 2003; Sourisseau et al., 2006; Georgiadis et al., 2010). Although it remains unclear whether DbpA regulates the apical junctional complex integrity, the mechanosensitive recruitment of DbpA may modify its transcriptional response and control the apical junctional complex integrity in a homeostatic manner.

### Roles of the actin cytoskeleton in epithelial junction homeostasis

The present findings suggest that actin polymerization plays a critical role in maintaining the junction integrity when TJs are perturbed. We further demonstrated that actin reassembly preceded junction reformation during the junction repair process, suggesting that the junction repair occurs in an actin filament-guided manner at least in claudin/JAM-A KO cells. Taken together with its well-established roles in junction assembly (Yonemura et al., 1995; Krendel and Bonder, 1999; Vasioukhin et al., 2000; Verma et al., 2004), these results suggest that the actin cytoskeleton plays pivotal roles in cell-cell junction repair and homeostasis. Consistent with this notion, it was reported that small breakages of TJs can be repaired by transient localized activations of Rho, termed Rho flares, that promote junction repair through actin polymerization and myosin II-dependent contraction (Stephenson et al., 2019). These observations suggest that actin polymerization plays a pivotal role in regulating epithelial junction homeostasis.

The circumferential actin bundles were extremely well developed in claudin/JAM-A KO cells, suggesting that the loss of TJs promotes the circumferential actin bundle development in a compensatory manner to maintain the apical junction integrity. Although the detailed signaling mechanism involved in the regulation of circumferential actin bundle formation in response to the loss of TJs remains to be fully clarified, it may involve the signaling pathways downstream of the TJ membrane proteins (Zihni et al., 2016), including the mechanosensitive calcium channels, guanine nucleotide exchange factors including p114RhoGEF and p115RhoGEF, and Shroom3 (Choi et al., 2016; Haas et al., 2020; Varadarajan et al., 2022; Chumki et al., 2022).

The present findings suggest that the ROCK-LIMK-cofilin pathway may regulate the mechanical resistance of apical junctions by promoting actin polymerization. Consistently, the RhoA-ROCK-LIMK-cofilin pathway was reported to promote actin assembly, resulting in stabilization of the apical junctional complex in colon carcinoma cells (Ito et al., 2017). Meanwhile, cofilin-dependent actin turnover was required to stabilize the newly formed junctions during the tissue tension-dependent junction remodeling in the *Drosophila* pupal wing (Ikawa et al., 2018), suggesting that the roles of cofilin may differ when the junctions are undergoing dynamic remodeling. ROCK-LIMK-cofilin-dependent turnover of the actin cytoskeleton may be involved in reorganization of the actin network to protect the junctions from catastrophic collapse when the junction integrity is challenged during morphogenesis.

In conclusion, we have demonstrated that TJ membrane proteins and the actin cytoskeleton coordinately regulate the apical junctional complex integrity when epithelial cells are mechanically challenged. Further analyses of the molecular mechanisms that regulate the coordinated response of cell junctions and the actin cytoskeleton to mechanical stress, as well as the quantitative measurements of the responses of cell junctions and the actin cytoskeleton to mechanical load, should deepen our understanding of epithelial homeostasis in the future.

## Materials and Methods

### Cell culture

MDCK II cells (Richardson et al, 1981) derived from the canine kidney were provided by Masayuki Murata (Tokyo Institute of Technology), and mouse L fibroblasts (L tk^–^) (Murayama-Okabayashi et al., 1971; Nagafuchi and Takeichi, 1987) were provided by Masatoshi Takeichi (RIKEN). The establishment of claudin/JAM-A KO cells was described previously (Otani et al; 2019). All cells were maintained in low-glucose DMEM (Nissui; #05919) supplemented with 2 mM L-glutamine (Nacalai Tesque; # 16948) and 10% FBS (Bio-West; #S1820-500) at 37°C under 5% CO_2_. The medium was changed every 2–3 d. ROCK inhibitor Y-27632 (Wako; # 030-24021) (Uehata et al., 1997) was used at 10 μM, myosin II inhibitor (-)-blebbistatin (Wako; #021-17041) (Straight et al., 2003) was used at 100 μM, LIMK inhibitor BMS-5 (Selleck; #S0185) (Ross-Macdonald et al., 2008) was used at 10 μM, and actin polymerization inhibitor latrunculin A (Calbiochem; #428021) (Coué et al., 1987) was used at 0.3 μM. The drugs were administered to confluent monolayers, and the cells were incubated for 1 h (latrunculin A) or 3 h (other inhibitors) before further analyses.

### Molecular biology

Canine ZO-1 was cloned from MDCK II cells using the following primers: 5′-TTGCTAGCATGTCCGCCAGAGCTGCGGC-3′, 5′-CCGTCGACAAAGTGGTCAATCAGGACAGAAACACAGTT-3′. Superscript IV reverse transcriptase (Invitrogen; #18090010) was used for cDNA synthesis, and cloning was performed by using KOD-Plus-Neo (TOYOBO; #KOD-401). Amplified cDNA was cloned into pEGFP-N3, pCANw-Sal-EGFP (Ichii and Takeichi, 2007), or pCAH-Sal-EGFP, wherein the neomycin-resistance gene of pCANw-Sal-EGFP was replaced with a hygromycin resistance gene, using an In-Fusion HD Cloning Kit (Takara; #Z9649N). pCANw-HA-cZO1-Flag was created by inserting an HA tag into the N-terminus of cZO-1 by PCR using the following primers: HA-forward: 5′-ATGTATCCGTATGATGTTCCGGATTATGCAAAGCTTATGTCCGCCAGAGCTG CGGCCG-3′; and Flag-reverse: 5′-TCGTCGTCCTTGTAGTCGACAAAGTGGTCAATCAGGACAGAAACACAGTTTGCTCCAACAAGG-3′. The PCR product was inserted into pCANw-Sal-Flag, wherein the GFP tag in pCANw-Sal-EGFP was replaced with a Flag tag. LifeAct7-mCherry was a gift from Michael Davidson (Addgene plasmid #54491) (Riedl et al., 2008), and cloned to pCAB-Sal, wherein the neomycin-resistance gene in pCANw was replaced with a blasticidin S-resistance gene.

Mouse claudin-1 (Furuse et al., 1998) and human JAM-A (Itoh et al., 2001) were cloned into pCANw. The mutant constructs were generated by PCR using KOD Plus ver.2 (TOYOBO; #KOD-211) and the following primer sets: Cl1-F147A-forward: 5′-TCAAGAAGCCTATGACCCCTTGACC-3′ and Cl1-F147A-reverse: 5′-GGTCATAGGCTTCTTGAACAATTCTGTTTCC-3′; Cl1-ΔYV-forward: 5′-GGGAAAGACTGACTCGAGTACAAGG-3′ and Cl1ΔYV-reverse: 5′-CTCGAGTCAGTCTTTCCCACTAGAAGG-3′, JAMA-ΔDL1-forward: 5′-GGCATTGGGCCTCATCGTGCTTGTGCCTCCATCCAAGCCTACAGTTAAC-3′ and JAMA-ΔDL1-reverse: 5′-GCACGATGAGGCCCAATGCCAGGGAGCACAACAGGATC-3′; JAMA-ΔLV-forward: 5′-ACCTCAAGCTTCTGAGTCGACTACAAGGACGACG-3′ and JAMA-ΔLV-reverse: 5′-CGACTCAGAAGCTTGAGGTCTGTTTGAATTCTCC-3′.

### Genome editing and transfection

Claudin/JAM-A/occludin KO cells and claudin/JAM-A/CAR KO cells were generated by introducing Cas9-gRNA RNP complexes into claudin/JAM-A KO cells. CRISPR RNA and trans-activating CRISPR RNA were synthesized by IDT and annealed, and subsequently complexed with Cas9 (IDT) to form the Cas9-gRNA RNP complex. Next, 100 pmol of Cas9 and 120 pmol of gRNA duplex were introduced into 1×10^5^ to 1×10^6^ cells using a CUY21 Pro-Vitro electroporator (Nepagene) under the following conditions: prepulse, 150 V for 10 ms; postpulses, 10× 20 V, 50 ms pulses at 50 ms intervals. The target sequences of the gRNAs were as follows (PAM sequences are underlined): occludin: 5′-GCACCGAGCAATGATGTGTA<u>CGG</u>-3′; and CAR: 5′-ACCCTTAGTCCAGAAGACCA<u>GGG</u>-3′. Electroporated cells were screened by immunofluorescence, and multiple KO cell clones were isolated and cultured. KO was verified by immunofluorescence, western blotting, and genomic sequencing.

Stable transfectants were isolated after transfection of the expression vectors using Lipofectamine® LTX reagent with PLUS™ reagent (Thermo Fisher Scientific; #A12621) in accordance with the manufacturer’s instructions. Transfected cells were selected using 500 μg/ml G418 (Nacalai Tesque; #16512-26), 200 μg/ml hygromycin B (Nacalai Tesque; #09287-84), or 2 μg/ml blasticidin S (Wako; #029-18701). Multiple surviving clones were isolated, and the expression of each transgene was confirmed by immunofluorescence and western blotting. At least three clones were isolated for each construct.

### Antibodies

The following primary antibodies were used for immunohistochemistry and western blotting analyses: mouse monoclonal anti-ZO-1 (clone T-8754) (Itoh et al., 1991); rabbit polyclonal anti-ZO-1 (Thermo Fisher Scientific; #61-7300); rabbit polyclonal anti-ZO-2 (Thermo Fisher Scientific; #38-9100); rabbit polyclonal anti-claudin-1 (Thermo Fisher Scientific; #51-9000); rabbit polyclonal anti-human JAM-A (Invitrogen; #PA5-120157); rabbit polyclonal anti-CAR (kindly provided by Jeffrey M. Bergelson) (Cohen et al., 2001); rat monoclonal anti-occludin (clone MOc37) (Saitou et al, 1997); rabbit polyclonal anti-occludin (Saitou et al, 1997); rat monoclonal E-cadherin (clone ECCD-2; kindly provided by Masatoshi Takeichi) (Shirayoshi et al., 1986); rabbit polyclonal anti-1/s-afadin (Sigma-Aldrich; #A0224); rabbit polyclonal anti-α-catenin (Sigma-Aldrich; #C2081); rat monoclonal α-18, which recognizes the extended conformation of α-catenin (kindly provided by Akira Nagafuchi) (Nagafuchi and Tsukita, 1994; Yonemura et al., 2010); rabbit polyclonal anti-MHCIIA (Sigma-Aldrich; #M8064); rabbit polyclonal anti-MHCIIB (BioLegend; #909901); mouse monoclonal α-tubulin (clone DM1A; Invitrogen; #14-4502-82) (Blose et al., 1984); rat monoclonal anti-HA (clone 3F10; Roche); and mouse monoclonal anti-FLAG (Wako; #014-22383).

The following secondary antibodies were used: Alexa Fluor 488-conjugated donkey anti-mouse IgG (Molecular Probes; #A21202); Alexa Fluor 488-conjugated donkey anti-rabbit IgG (Molecular Probes; #A21206); Alexa Fluor 488-conjugated donkey anti-rat IgG (Molecular Probes; #A21208), Alexa Fluor 555-conjugated donkey anti-mouse IgG (Invitrogen; #A31570); Alexa Fluor 555-conjugated donkey anti-rabbit IgG (Invitrogen; #A31572); Alexa Fluor 555-conjugated goat anti-rat IgG (Invitrogen; #A21434); Alexa Fluor 647-conjugated donkey anti-mouse IgG (Invitrogen; #A31571); Alexa Fluor 647-conjugated phalloidin (Invitrogen; #A22287); sheep anti-mouse IgG HRP-conjugated whole antibody (Cytiva; #NA931V); donkey anti-rabbit HRP-conjugated F(a’b’)2 fragment (Cytiva; #NA9340V); and anti-rat IgG HRP-conjugated antibody (Cytiva; #NA935V).

### Immunofluorescence

For immunofluorescence of MDCK II cells and their derivatives, 1×10^5^ cells were cultured on Transwell polycarbonate filters (0.4 µm pore size; Corning; #3401) for 5–7 d. For immunofluorescence of L cells, 1.5×10^5^ cells were plated on coverslips in 35 mm dishes and cultured for 2 d. Cells were fixed with 1%–2% PFA (RT,15 min), 10% TCA (4°C, 15 min), or 100% methanol (–20°C, 10 min). Specifically, 1% PFA fixation was used for anti-α-catenin, α-18, anti-vinculin, anti-E-cadherin, anti-afadin, anti-HA, and anti-Flag, 2% PFA fixation was used for anti-ZO-1, anti-JAM-A, phalloidin, anti-myosin IIA and anti-myosin IIB, TCA fixation was used for anti-ZO-2; and methanol fixation was used for anti-occludin. After fixation, the cells were washed three times with PBS, permeabilized with 0.1% TritonX-100 in PBS (RT, 15 min), rinsed once with PBS, and blocked with 10% FBS (RT, 30 min). For cells cultured on Transwell filters, the filters were excised using scalpels. All cells were incubated with the primary antibodies diluted in blocking solution (RT, 1 h), washed three times with PBS, and incubated with secondary antibodies diluted in blocking solution (RT, 1 h). Finally, the cells were washed three times with PBS and mounted using FluoroSave™ Reagent (Calbiochem; #345789). For STED imaging, samples were mounted using ProLong™ Diamond Antifade Mountant (Thermo Fisher Scientific; #P36961) and covered with coverslips (Matsunami; #No.1S HT; 0.17±0.005 mm).

### Confocal imaging

Confocal images were obtained using an AX R confocal laser scanning microscope system mounted on an Eclipse Ti2 inverted microscope with a CFI PLAN Apochromat lambda D 60× oil (NA 1.42) objective and diode lasers (488/561/640 nm) accompanied by the NIS-Elements C Imaging software (all from Nikon Solutions).

For STED imaging, cells were imaged using a TCS SP8 STED confocal system mounted on a DMI8 inverted microscope with an HC PL APO CS2 100× oil (NA 1.4) objective and diode lasers (confocal: 488/555 nm; STED: 592/660 nm) accompanied by Application Suite X acquisition software (all from Leica Microsystems).

### Time-lapse imaging

For time-lapse imaging, 1×10^5^ cells were plated at the center of glass-based dishes (Iwaki; #3910-035). After attachment, the cells were cultured in low glucose, phenol red-free DMEM (Nacalai Tesque; #08490-05), supplemented with 10% FBS and penicillin-streptomycin-glutamine (Gibco; #10378-016). Time-lapse imaging was performed using a CellVoyager CV1000 spinning disc confocal imaging system (Yokogawa) using 40× dry (NA 1.3) or 60× Silicone (NA 1.35) objective lenses with diode lasers (488/561 nm). The cells were imaged at 1 min intervals for 10 h (for ZO-1-GFP/LifeAct7-mCherry two-color imaging) with z-sectioning to 22 µm and 0.6 µm step size, or at 12 min intervals for 5 d (ZO-1-GFP movies) with z-sectioning to 22 µm and 1 µm step size. Movie acquisition was performed using CV1000 software (Yokogawa).

### Quantitative image analyses

All images were analyzed and processed using Fiji/ImageJ 1.53t software (National Institutes of Health). Brightness and contrast adjustments without any nonlinear adjustments were applied for qualitative presentation of images when necessary.

Junction breakage was quantified as follows. First, noise reduction was performed by application of band-pass and median filters, and the junction outlines were extracted by thresholding and binarization. After removal of outliers and manual removal of misannotated junctions, the image was skeletonized. Next, an averaging filter was applied to the skeletonized image, and the endpoints were extracted by thresholding and subsequent application of a band-pass filter to remove the edges of the bicellular junctions. The number of endpoints was counted using the “analyze particles” function, and the area of the region of interest was measured. Finally, the number of endpoints was divided by the area to yield the number of endpoints per unit area.

The degree of cell stretching was quantified by measuring the outline of the cell of interest. The intensity of ZO-1-GFP was measured by measuring the intensity along the outline of the cell. The beginning of a stretching event was defined as the starting point, yielding L_0_ and I_0_, with ΔL and ΔI defined as the maximum relative length change and the maximum relative intensity change during a single stretching event, respectively. The degree of cell stretching was defined as 100×ΔL/ L_0_ (%), and the change in ZO1-GFP intensity was defined as 100×ΔI/ I_0_ (%).

The distance between the N- and C-termini of ZO-1 was measured by performing a line scan across pairs of HA- and Flag-tag puncta. Only pairs that appeared as doublets were measured. The distance between the fluorescence peaks of the HA-and Flag-tags was measured. The order parameter was quantified as follows. First, the angle between the N- and C-termini of ZO-1 was measured. Next, the mean angle of the particles within a single image was defined, and the deviation of individual vectors from the average vector was measured to determine the relative angle θ. Finally, the order parameter *S* = (3cos^2^θ-1)/2 was calculated for each vector.

Graph generation and statistical tests (*t*-test, unpaired, two-tailed) were performed using Excel software (Microsoft) or R (R foundation).

### Western blotting

Cells grown on 35 mm dishes were rinsed once with ice-cold PBS, lysed with 200 µL of Laemmli sample buffer supplemented with 100 mM DTT, sonicated, and boiled at 97°C for 5 min. Samples were separated by SDS-PAGE using standard methods, with 20 µl of sample loaded per lane. The separated proteins were transferred to 0.45 μm pore Protran nitrocellulose transfer membranes (Whatman; #10-401-196). The membranes were blocked with 5% skim milk in TBS/0.1% Tween-20 for 1 h at RT, and incubated with primary antibodies diluted in the blocking solution overnight at 4°C. The membranes were rinsed three times with TBS/0.1% Tween-20 and incubated with secondary antibodies diluted in blocking solution for 1 h at RT. After three washes with TBS/0.1% Tween-20, signals were detected by chemiluminescence using ECL Prime Western Blotting Detection Reagents (Cytiva; #RPN2232). Images were obtained using an LAS3000 mini system (Fujifilm), and data were processed and analyzed using Fiji/ImageJ 1.53t software.

### Online supplemental material

Fig. S1 shows the actomyosin organization in claudin/JAM-A KO cells. Fig. S2 shows the characterization of claudin-1 and JAM-A mutants. Fig. S3 shows the verification of claudin/JAM-A/occludin and claudin/JAM-A/CAR KO cells.

Video 1 shows the dynamics of ZO-1-GFP in MDCK II cells. Video 2 shows the junction breakage events in claudin/JAM-A KO cells expressing ZO-1-GFP. Video 3 shows the actin dynamics during junction breakage observed by two-color time-lapse imaging of claudin/JAM-A KO cells expressing ZO-1-GFP and LifeAct-mCherry. Video 4 shows the actin dynamics during junction repair observed by two-color time-lapse imaging of claudin/JAM-A KO cells expressing ZO-1-GFP and LifeAct-mCherry.

## Data availability

The data underlying this study are available from the corresponding authors upon reasonable request.

## Acknowledgments

We thank Akira Nagafuchi, Masayuki Murata, Jeffrey M. Bergelson, Masatoshi Takeichi, Michael Davidson, and the Developmental Studies Hybridoma Bank for kindly providing reagents; Mika Watanabe and Yuichiro Kano for technical assistance; Motohiro Nishida for allowing us to use the microplate reader; and all members of the Furuse laboratory for discussions and comments. We also thank Alison Sherwin from Edanz for editing a draft of this manuscript.

This work was supported by a JSPS Grant-in-Aid for Challenging Exploratory Research (16K15226, to MF), JSPS Grants-in-Aid for Scientific Research (B) (26291043, 18H02440, 21H02523, to MF), JSPS Grant-in-Aid for Scientific Research (C) (18K06234, TO), JSPS Grants-in-Aid for Young Scientists (B) (16K18544, to TO; 21K15095, to SF), MEXT/JSPS Grant-in-Aid for Scientific Research on Innovative Areas (17H05627, to TO), MEXT/JSPS Grant-in-Aid for Transformative Research Areas (21H05286, to TO), JST PRESTO (JPMJPR204, to TO); MEXT/JSPS Grant-in-Aid for Scientific Research on Innovative Areas – Platforms for Advanced Technologies and Research Resources “Advanced Bioimaging Support”; (JP16H06280, JP22H04926), the Inamori Foundation (to TO), The Japan Spina Bifida & Hydrocephalus Research Foundation (to TO), and the Takeda Science Foundation (to MF, TO).

The authors declare no competing financial interests.

## Author contributions

Conceptualization, Funding acquisition, Supervision, Project administration: T. Otani and M. Furuse; Resources: T. Otani, T.P. Nguyen, M. Tsutsumi, S. Fujiwara, T. Nemoto, T. Fujimori, and M. Furuse; Formal analysis: T. Otani and T.P. Nguyen; Investigation, Visualization: T.P. Nguyen and T. Otani; Writing – original draft: T. Otani, T.P. Nguyen, and M. Furuse; Writing – review & editing: all authors.

## Abbreviations List

AJ: adherens junction
CAR: coxsackie and adenovirus receptor
JAM: junctional adhesion molecule
KO: knockout
STED: stimulated-emission depletion
TJ: tight junction

## Legends for Supplemental Figures

**Figure S1. Actomyosin organization in claudin/JAM-A KO cells.**

(A–C) F-actin/myosin IIA/ZO-1 triple staining in MDCK II cells. (A’ –C’) F-actin/myosin IIA/ZO-1 triple staining in claudin/JAM-A KO cells. It should be noted that the circumferential actin bundles were extensively developed. (D, E) Myosin IIB/ZO-1 double staining in MDCK II cells. (D’, E’) Myosin IIB/ZO-1 double staining in claudin/JAM-A KO cells. Myosin IIB was strongly localized to the extensively developed circumferential actin bundles. (F–H) α-18/α-catenin/ZO-1 triple staining in MDCK II cells. The α-18 antibody recognizes a cryptic epitope exposed by the tension-dependent stretching of α-catenin. Weak staining of α-18 along the apical cell junctions was observed. (F’–H’) α-18/α-catenin/ZO-1 triple staining in claudin/JAM-A KO cells. Strong staining of α-18 was observed at the cell-cell junctions, while weak staining was present at the junction breakage sites. (I, J) Vinculin/ZO-1 double staining in MDCK II cells. Junctional staining was hardly visible in confluent monolayers. (I’, J’) Vinculin/ZO-1 double staining in claudin/JAM-A KO cells. Strong accumulation of vinculin was observed at the apical cell junctions, while weak staining was noted at the junction breakage sites. Scale bar: 10 μm.

**Figure S2. Characterization of claudin-1 and JAM-A mutants.**

(A–C) Localization of claudin-1 and its mutants in L cells. Claudin-1 and its mutants were ectopically expressed in L cells, which lack endogenous cell-cell adhesion activity. Claudin-1 full-length (A) and claudin-1 [ΔYV] (C) were localized at cell-cell contacts (arrows), while claudin-1[F147A] (B) was not concentrated at cell-cell contacts. (D–F) Localization of JAM-A and its mutants in L cells. JAM-A full-length (D) and JAM-A[ΔLV] (F) were localized at cell-cell contacts (arrows), while JAM-A[ΔDL1] (E) was not localize at cell-cell contacts. (G–I) Double staining of claudin-1[FL] and ZO-1 in L cells. Claudin-1[FL] was able to recruit ZO-1 to cell-cell contacts (arrows). (J–L) Double staining of claudin-1[ΔYV] and ZO-1. Claudin-1[ΔYV] did not recruit ZO-1 to cell-cell contacts (arrowheads). (M–O) Double staining of JAM-A[FL] and ZO-1 in L cells. JAM-A[FL] was able to recruit ZO-1 to cell-cell contacts (arrows). (P–R) Double staining of JAM-A[ΔLV] and ZO-1. JAM-A[ΔLV] did not recruit ZO-1 to cell-cell contacts (arrowheads). Scale bars: 10 μm.

**Figure S3. Generation of claudin/JAM-A/occludin KO and claudin/JAM-A/CAR KO cells.**

(A) Western blotting of occludin, CAR, and α-tubulin in MDCK II, claudin/JAM-A KO, claudin/JAM-A/occludin KO, and claudin/JAM-A/CAR KO (clones 1 and 2) cells. Occludin expression was specifically lost in claudin/JAM-A/occludin KO cells, while CAR expression was specifically lost in claudin/JAM-A/CAR KO cells.

## Legends for Supplemental Videos

**Video 1. ZO-1-GFP dynamics in MDCK II cells.**

Time-lapse imaging of ZO-1-GFP expressed in MDCK II cells. The cells pushed and pulled one another, but the junction continuity was maintained. Maximum intensity projections of all z-sections are shown. Fig. 2A and 2B are cropped images from this movie.

**Video 2. Spontaneous junction breakage in claudin/JAM-A KO cells.**

Time-lapse imaging of ZO-1-GFP expressed in claudin/JAM-A KO cells. The cell junctions underwent spontaneous breakage when the cells were acutely stretched. A cell in the upper right subsequently underwent cytokinesis and junction fragmentation occurred. Maximum intensity projections of all z-sections are shown. Fig. 2C and 2D are cropped images from this movie.

**Video 3. Actin dynamics during junction breakage in claudin/JAM-A KO cells.**

Time-lapse imaging of ZO-1-GFP and LifeAct-mCherry expressed in claudin/JAM-A KO cells. The cell junctions-associated actin bundles were loosened and became detached from the cell junctions simultaneously with junction breakage. Maximum intensity projections of all z-sections are shown. Left: merged image; center: ZO-1-GFP; right: LifeAct-mCherry.

**Video 4. Actin dynamics during junction recovery in claudin/JAM-A KO cells.**

Time-lapse imaging of ZO-1-GFP and LifeAct-mCherry expressed in claudin/JAM-A KO cells. The cell junction-associated actin bundles reformed before the junction repair occurred. Maximum intensity projections of all z-sections are shown. Left: merged image; center: ZO-1-GFP; right: LifeAct-mCherry.

